## Supporting Information for "Development of a p62 biodegrader for autophagy targeted degradation"

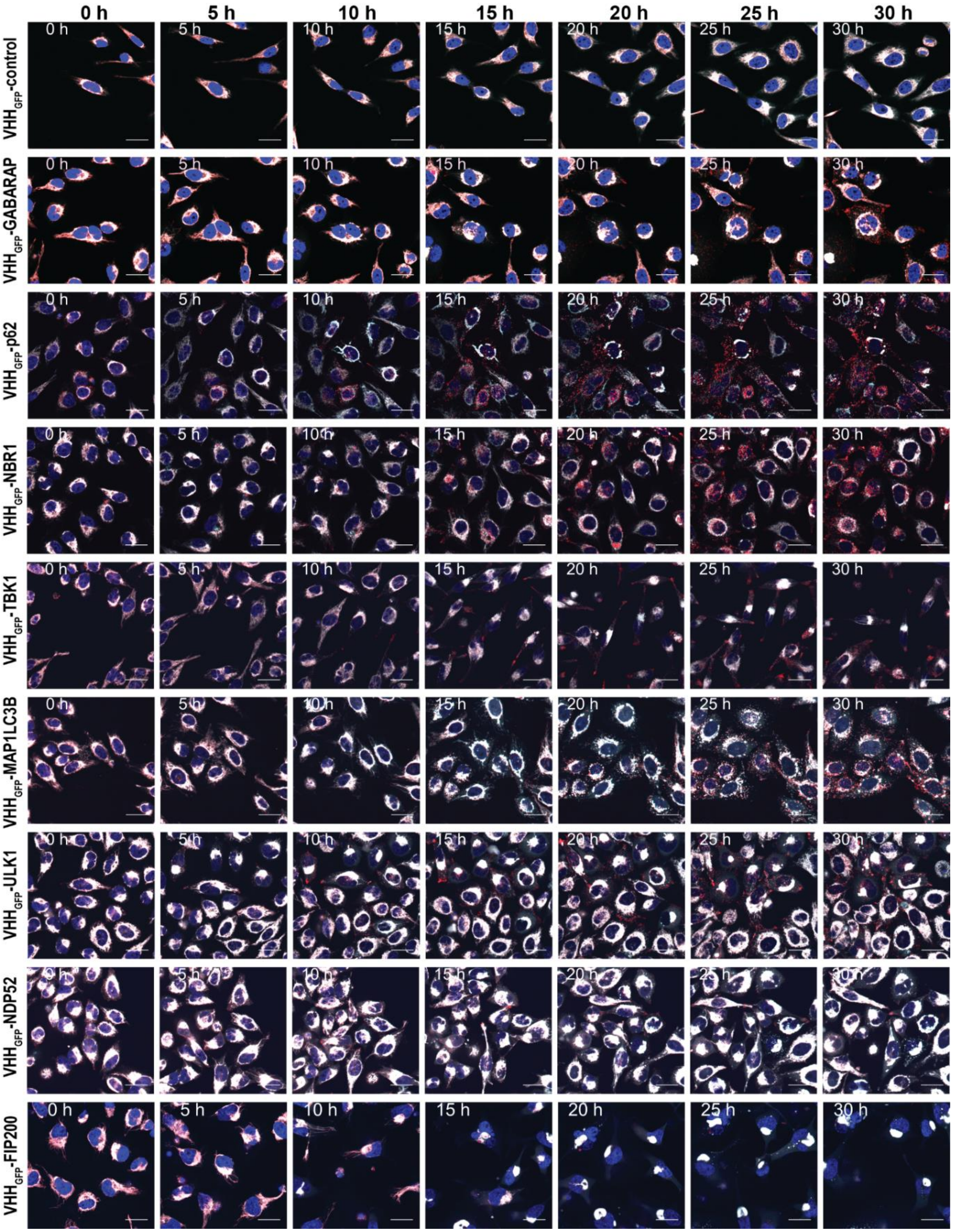


**Figure S1. Time-series imaging of HeLa mito-mCh-GFP cells expressing selected fusions of VHH$_{GFP}$ and autophagy effectors.**

Expression of the VHH$_{GFP}$ - autophagy effector fusion constructs was induced by addition of 1 µg mL$^{-1}$ doxycycline and the cells were immediately imaged (0h). Fluorescence images of the same field of views were taken every hour for 30 h. Red = mCh, cyan = GFP, blue = nuclei. Scale bars are 30 µm

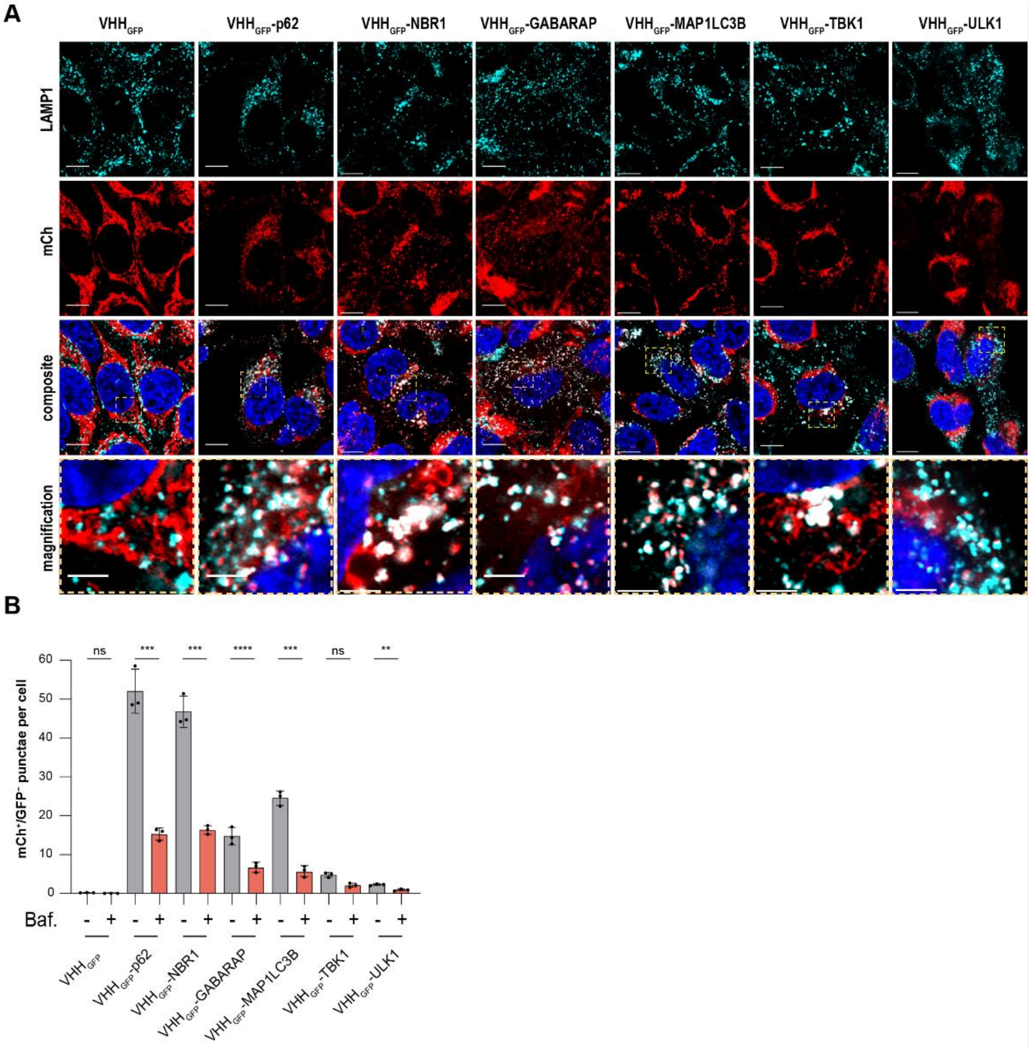


**Figure S2. Induced proximity of autophagy effectors results in lysosomal delivery of mitochondria in HeLa mito-mCh-GFP cells and is dependent on autophagic activity.**

(A) Representative immunofluorescence Airyscan images of HeLa mito-mCh-GFP cells expressing the fusions of $VHH_{GFP}$ and autophagy effectors for 72 h. Cells were fixed and immunostained. Red = mCh, cyan = LAMP1, blue = nuclei. Scale bars are 20 μm and 5 μm (magnification).

(B) Formation of $mCh^{+}/GFP^{-}$ punctae in HeLa mito-mCh-GFP cells expressing the fusions of $VHH_{GFP}$ and autophagy effectors for 72 h and treated with Bafilomycin A1 (100 nM) for 15 h. The data are shown as mean and standard deviation from 3 independent biological replicates. Statistical analysis was performed using an unpaired *t*-test comparing untreated with treated samples. ns = ($P > 0.05$); * = ($P \leq 0.05$); ** = ($P \leq 0.01$); *** = ($P \leq 0.001$); **** = ($P \leq 0.0001$)

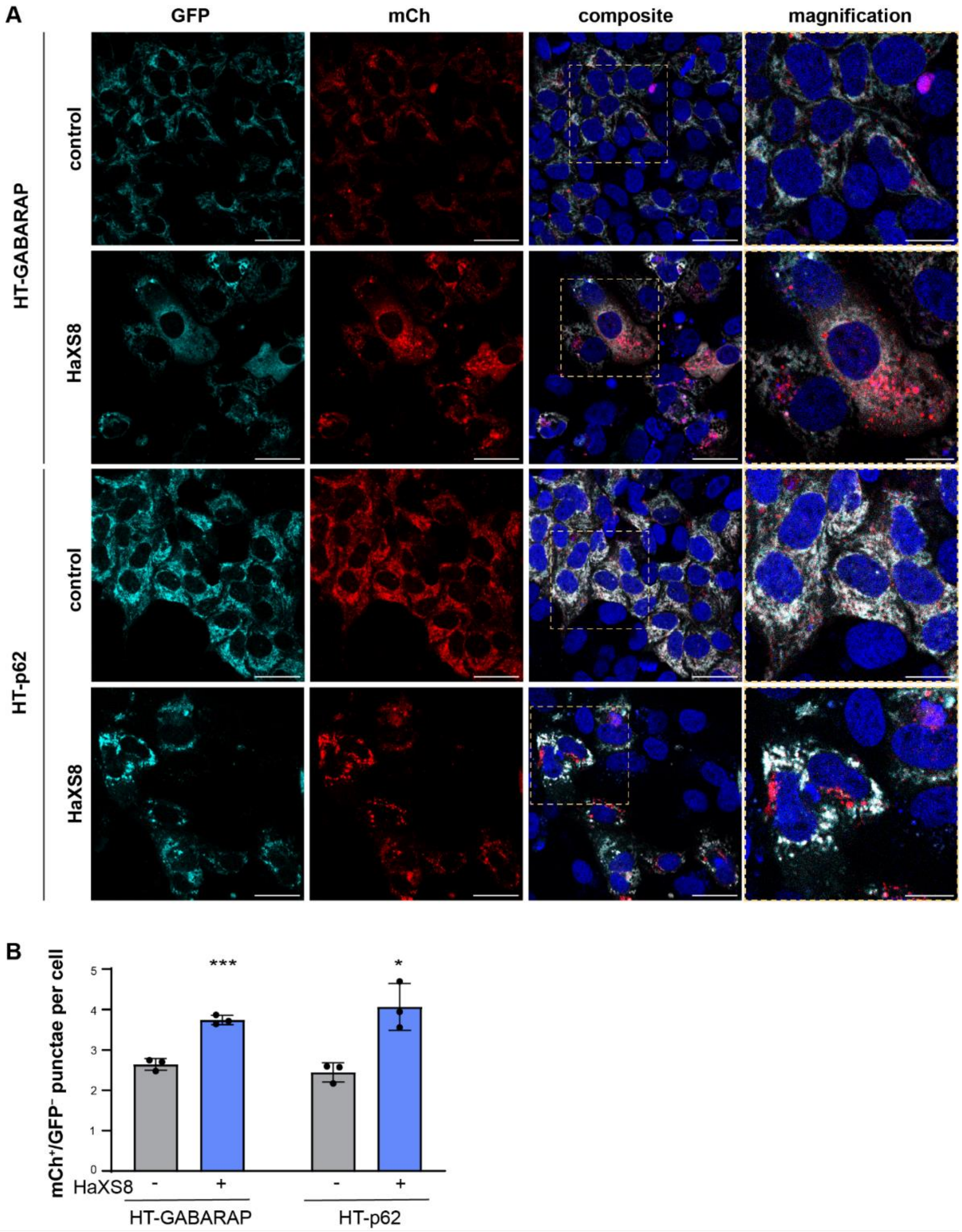


**Figure S3. Lysosomal delivery of mitochondria in Huh7 cells using HaXS8.**

(A) Confocal fluorescence microscopy images showing live Huh7 mito-mCh-GFP-SNAPtag cells expressing HaloTag-GABARAP or HaloTag-p62. Cells were treated with 100 nM HaXS8 for 72 h. Red = mCh, cyan = GFP, blue = nuclei. Scale bars are 30 µm and 15 µm (magnification).

(B) Quantification of the images displayed in panel (A). The data are shown as mean and standard deviation from n = 3 technical replicates. Statistical analysis was performed using parametric unpaired *t*-tests comparing untreated and treated samples. * = ($P < 0.05$); *** = ($P < 0.001$)

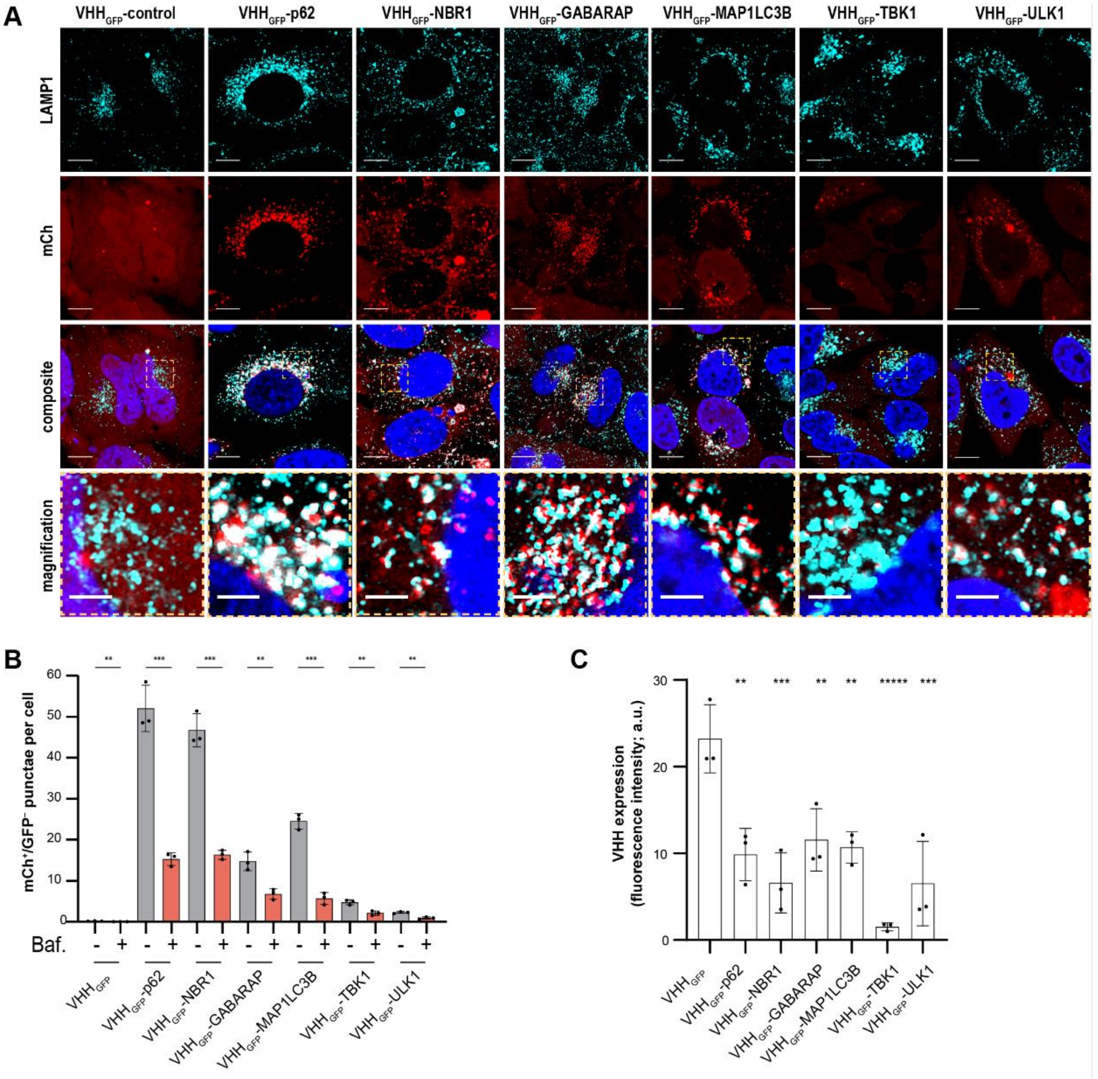


**Figure S4. Induced proximity of autophagy effectors results in lysosomal delivery of a soluble protein in HeLa cyt-mCh-GFP cells and is dependent on autophagic activity.**

(A) Representative immunofluorescence Airyscan images of HeLa mito-mCh-GFP cells expressing fusions of $VHH_{GFP}$ and autophagy effectors for 72 h. Cells were fixed and immunostained. Red = mCh, cyan = LAMP1, blue = nuclei. Scale bars are 20 µm and 5 µm (magnification).

(B) Formation of $mCh^+/GFP^-$ punctae in HeLa cyt-mCh-GFP cells expressing fusions of $VHH_{GFP}$ and autophagy effectors for 72 h and treated with Bafilomycin A1 (100 nM) for 15 h. The data are shown as mean and standard deviation from 3 independent biological replicates. Statistical analysis was performed using an unpaired *t*-test comparing untreated with treated samples. ns = ($P > 0.05$); * = ($P \leq 0.05$); ** = ($P \leq 0.01$); *** = ($P \leq 0.001$); **** = ($P \leq 0.0001$)

(C) Normalized expression levels of VHH in HeLa cyt-mCh-GFP cells 72 h post-induction with doxycycline. Fluorescence intensities were assessed by immunofluorescence microscopy and were normalized to the background fluorescence measured in untransfected HeLa mCh-GFP cells. The data are shown as mean and standard deviation from 3 independent biological replicates. Statistical analysis was performed using an ordinary one-way ANOVA with multiple comparison of each data point against untransfected cells. ns = ($P > 0.05$); * = ($P \leq 0.05$); ** = ($P \leq 0.01$); *** = ($P \leq 0.001$); **** = ($P \leq 0.0001$)

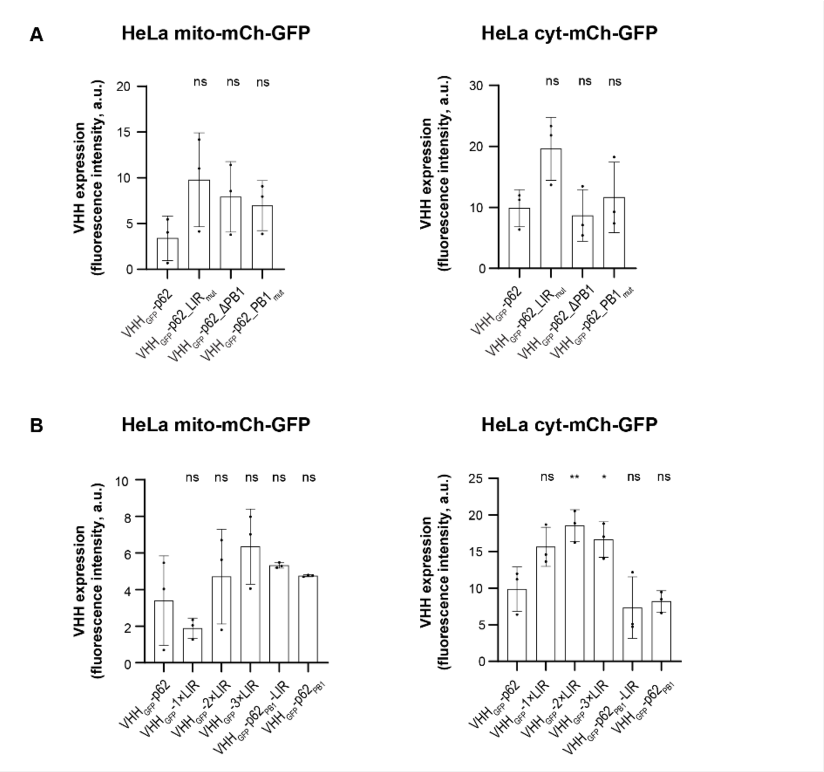


**Figure S5. Expression levels of $VHH_{GFP}$-p62 mutant fusion constructs.**

(A) Expression levels of $VHH_{GFP}$ fusions of p62 mutants in HeLa mito-mCh-GFP and HeLa cyt-mCh-GFP cells 72 h post-induction with doxycycline. Fluorescence intensities were assessed by immunofluorescence microscopy. The data are shown as mean and standard deviation from 3 independent biological replicates. Statistical analysis was performed using an ordinary one-way ANOVA with multiple comparison of each data point against $VHH_{GFP}$-p62. ns = ($P > 0.05$); * = ($P \leq 0.05$); ** = ($P \leq 0.01$); *** = ($P \leq 0.001$); **** = ($P \leq 0.0001$).

(B) Expression levels of $VHH_{GFP}$ fusions of 1-3 repeats of the p62 LIR peptide, the minimal degron PB1-LIR and the PB1 domain of p62 in HeLa mito-mCh-GFP and HeLa mCh-GFP cells 72 h post-induction with doxycycline. Fluorescence intensities were assessed by immunofluorescence microscopy. The data are shown as mean and standard deviation from 3 independent biological replicates. Statistical analysis was performed using an ordinary one-way ANOVA with multiple comparison of each data point against $VHH_{GFP}$-p62. ns = ($P > 0.05$); * = ($P \leq 0.05$); ** = ($P \leq 0.01$); *** = ($P \leq 0.001$); **** = ($P \leq 0.0001$).

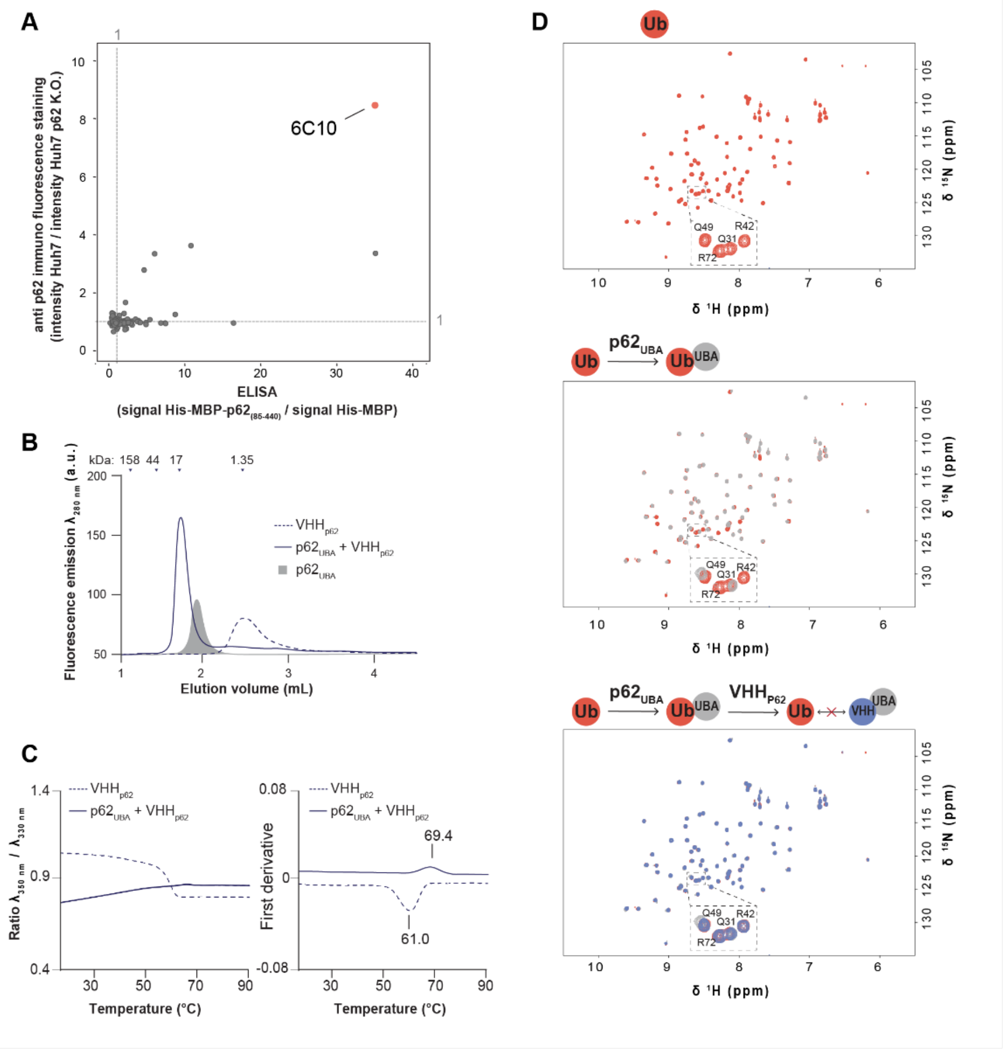


**Figure S6. Generation and analytical QC of VHH$_{p62}$.**

(A) Selection of the anti-p62 VHH with the lowest signal to noise ratio using immunofluorescence microscopy and ELISA assays. Immunofluorescence microscopy experiments were performed using periplasmic extracts of *E. coli* expressing VHHs followed by staining against VHH. The ratio of the fluorescence intensities obtained in Huh7 and Huh7 p62. K.O. was then calculated. ELISA assays were performed using periplasmic extracts of *E. coli* expressing VHHs. The ratio was calculated from the signal obtained from plates coated with the full antigen His-MBP-p62$_{(85\text{-}440)}$ or the control His-MBP.

(B) Size exclusion chromatography (SEC) of recombinant VHH$_{p62}$, p62$_{UBA}$, and a 1:1.5 mixture of VHH$_{p62}$ and p62$_{UBA}$.

(C) Nano differential scanning fluorimetry of VHH$_{p62}$ alone and a 1:2 mixture of VHH$_{p62}$ and p62$_{UBA}$.

(D) ($^{1}$H,$^{15}$N)-SOFAST-HSQC spectra of $^{13}$C,$^{15}$N-labeled ubiquitin, a 1:1 mixture of $^{13}$C,$^{15}$N-labeled ubiquitin and p62$_{UBA}$, and a 1:1:1.2 mixture of $^{13}$C,$^{15}$N-labeled ubiquitin, p62$_{UBA}$, and VHH$_{p62}$.

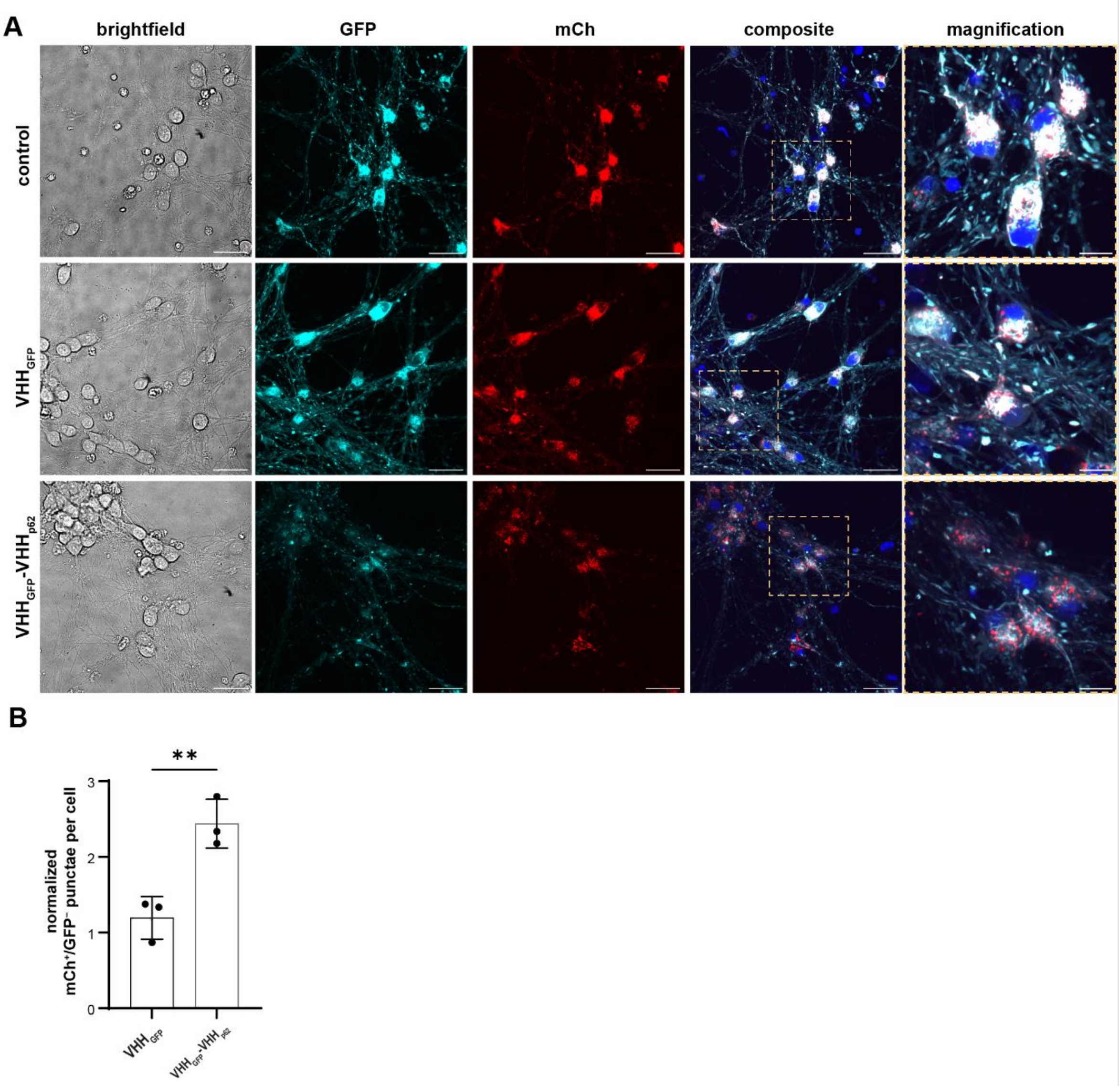


**Figure S7. Targeted degradation of mitochondria in iPSC derived NGN2 neurons.**

(A) Representative confocal fluorescence microscopy images showing live iPSC-derived NGN2 neurons expressing mito-mCh-GFP and $VHH_{GFP}$, $VHH_{GFP}$-GABARAP, $VHH_{GFP}$-p62, or $VHH_{GFP}$-$VHH_{p62}$ for 7 days. Red = mCh, cyan = GFP, blue = nuclei. Scale bars are 20 µm

(B) Quantification of the images displayed in panel (A). The data are shown as mean from 2 independent biological replicates. Statistical analysis was performed using an ordinary one-way ANOVA with multiple comparison of each data point against $VHH_{GFP}$. ns = ($P > 0.05$); * = ($P \leq 0.05$); ** = ($P \leq 0.01$); *** = ($P \leq 0.001$); **** = ($P \leq 0.0001$).

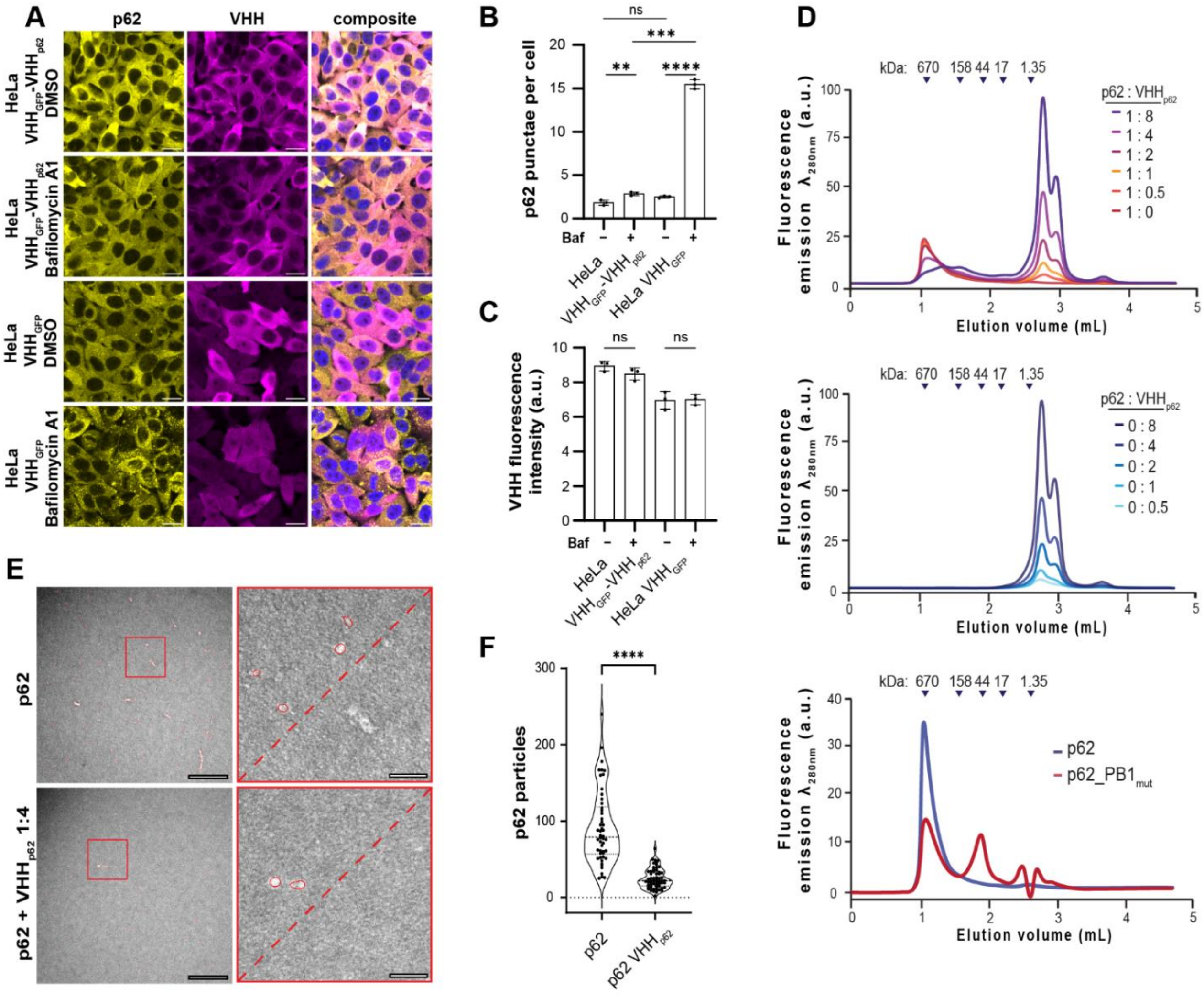


**Figure S8. Binding of $VHH_{p62}$ to p62 inhibits p62 self-oligomerization.**

(A) Representative images of HeLa cells expressing $VHH_{GFP}$ or $VHH_{GFP}$-$VHH_{p62}$ for 24 h and treated with DMSO or Bafilomycin A1 (100 nM) for 4 h. Cells were fixed and immunostained. yellow = p62, magenta = VHH. Scale bars are 20 µm.

(B) Quantification of p62 punctae in HeLa cells expressing $VHH_{GFP}$-$VHH_{p62}$ or $VHH_{GFP}$ after treatment with Bafilomycin A1 (100 nM) for 4 h. The data are shown as mean and standard deviation from 3 biological replicates. Statistical analysis was performed using unpaired *t*-tests performed on untreated versus treated samples and between the different cell lines under basal conditions and after treatment with Bafilomycin A1. ** = ($P \leq 0.01$); *** = ($P \leq 0.001$); **** = ($P < 0.0001$).

(C) Expression levels of VHH in HeLa cells expressing $VHH_{GFP}$ or $VHH_{GFP}$-$VHH_{p62}$ 24 h post induction with doxycycline. Expression levels were assessed by immunofluorescence microscopy. The data are shown as mean and standard deviation from 3 biological replicates. Statistical analysis was performed using unpaired *t*-tests comparing treated and untreated cells. ns = ($P > 0.05$).

(D) SEC analysis of recombinant full-length p62 in the presence of increasing concentrations of recombinant $VHH_{p62}$.

(E) SEC analysis of increasing concentrations of recombinant $VHH_{p62}$.

(F) SEC analysis of recombinant wild type p62 and $p62\_PB1_{mut}$.

**(G)** Negative staining transmission electron microscopy images of recombinant full length p62 with and without addition of recombinant $VHH_{p62}$ (left) and quantification of high-molecular weight particles formed by p62 (right). Scale bars are 200 and 50 nm (magnification). The data are shown as individual data points, mean and standard deviation from from 3 technical replicates. Statistical analysis was performed using an unpaired *t*-test performed on untreated versus treated samples. **** = ($P < 0.0001$).
